# Pathological cortico-STN beta coupling in Parkinson’s disease is confined to beta bursts

**DOI:** 10.64898/2026.04.09.717378

**Authors:** Christopher A. Beaudoin, Andrew B. O’Keeffe, Jeong Woo Choi, Amirezza Alijanpourotaghsara, Martin J. Gillies, Ashwini A. Oswal, Nader Pouratian, Alexander L. Green

## Abstract

Abnormal beta-band activity (13-30 Hz) within the cortico-basal ganglia network is a hallmark of Parkinson’s disease (PD) and is closely linked to motor impairment. Pathological beta activity in the subthalamic nucleus (STN) occurs predominantly as brief, high-amplitude bursts rather than continuous oscillations. Although beta-band coherence between the STN and cortex increases during bursts, it remains unclear whether cortico-STN beta coupling persists outside these bursts. Using intraoperative STN local field potentials and simultaneous cortical electrocorticography from seven patients undergoing deep brain stimulation implantation surgery, cortico-STN beta coupling during burst and non-burst epochs was compared. Coupling was assessed using magnitude-squared coherence and the debiased weighted phase lag index (dwPLI) and compared against surrogate distributions generated by circular time-shifting. Both coupling metrics were significantly elevated during burst epochs relative to non-burst periods. During non-burst epochs, coupling collapsed to surrogate levels, indicating no evidence of sustained synchronization. Time-resolved analyses further demonstrated that elevated coupling was confined to burst epochs. Although a subset of motor cortical contacts exhibited elevated baseline coherence, coupling was less evident using dwPLI. These findings suggest that pathological cortico-STN beta coupling in PD is preferentially expressed during beta bursts rather than sustained across non-burst epochs, with implications for adaptive neuromodulation strategies.

## 1. Introduction

Abnormal beta-band activity (∼13-30 Hz) within the cortico-basal ganglia network is a hallmark of Parkinson’s disease (PD), associated with motor symptoms such as bradykinesia and rigidity[1–3]. Beta oscillations in the subthalamic nucleus (STN) and increased coherence between the STN and motor cortex have been linked to impaired motor function and are attenuated by dopaminergic therapy and deep brain stimulation[4,5]. Pathological STN beta activity occurs predominantly as brief, intermittent high-amplitude bursts rather than continuous rhythms[6]. The duration, rate, and timing of these beta bursts correlate more closely with motor impairment than average beta power[7,8]. Beta bursts may reflect transient increases in network synchrony rather than isolated events[9]. Consistent with this view, cortico-STN beta coherence increases during beta bursts and diminishes outside these events[10,11].

However, while beta activity is increasingly understood to be temporally structured, it remains unclear whether cortico-STN beta coupling is effectively absent during non-burst periods or whether weaker, tonic coupling persists outside bursts[12]. To address this, intraoperative recordings of STN local field potentials (LFPs) and simultaneous cortical electrocorticography (ECoG) were analyzed from seven patients undergoing deep brain stimulation implantation surgery for PD. Cortico-STN beta coupling was quantified separately during beta burst and non-burst epochs and compared against surrogate data to control for spurious correlations. This surrogate-controlled approach tested whether cortico-subcortical beta-band coupling is confined to burst epochs or persists outside them.

## 2. Results

### 2.1. Cortico-STN beta coupling during beta bursts is distinct from non-burst epochs

Given prior reports that high-amplitude beta bursts are associated with increased network coherence, whereas non-burst periods are not, the difference in cortico-STN beta coherence between burst and non-burst epochs was assessed[10,11]. Band-pass filtered (13-30 Hz) and trimmed STN LFP and cortical ECoG signals were analyzed using magnitude-squared coherence (MATLAB mscohere), with bursts defined from the smoothed Hilbert envelope of the STN beta signal (>75th percentile; non-burst <50th percentile; minimum duration 0.1 s). Across STN-cortical contact pairs (STN contacts 5-8; cortical contacts 9-16), beta coherence was significantly higher during bursts than non-burst periods (paired Wilcoxon signed-rank test, p = 5.37 × 10^-42^; paired t-test, p = 8.48 × 10^-76^), with a strongly positive distribution of burst-versus-non-burst differences (Figure 1A, C). To assess robustness to amplitude-dependent scaling, the same analysis was repeated using the debiased weighted phase lag index (dwPLI), a phase-based metric insensitive to zero-lag and magnitude effects. Burst dwPLI was significantly greater than non-burst dwPLI (paired Wilcoxon signed-rank test, p = 1.98 × 10^-42^; paired t-test, p = 7.74 × 10^-121^), with a positive distribution of burst-versus-non-burst differences (Figure 1B, D). These effects remained significant in linear mixed-effects models with random intercepts for patient and electrode contact. Together, these findings demonstrate that cortico-STN beta coupling is elevated during burst epochs relative to non-burst periods across both magnitude- and phase-based metrics.

**Figure 1.**
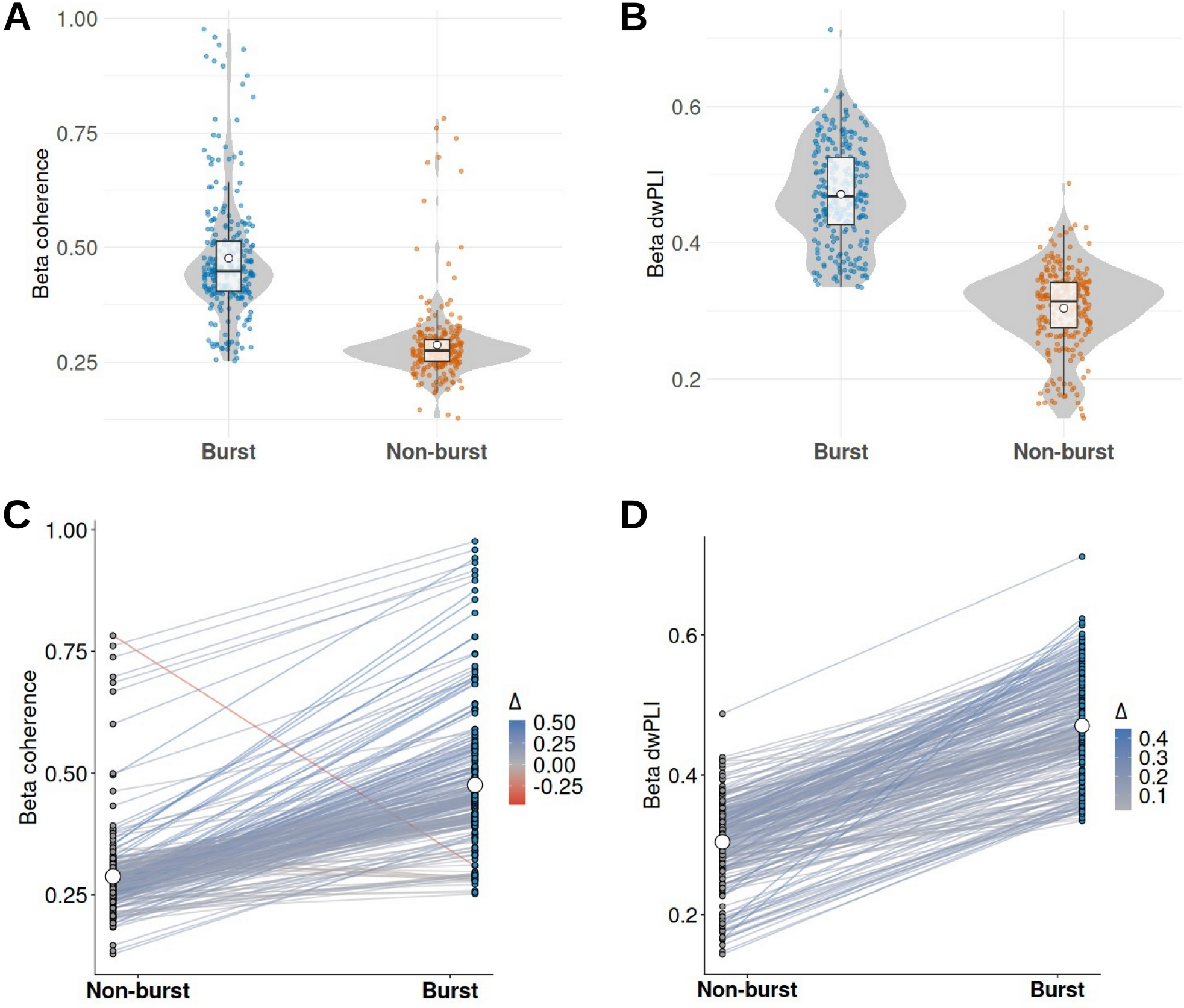
Cortico-STN beta coupling during burst and non-burst epochs Mean beta coherence (A) and dwPLI (B) for each STN-cortical contact pair during burst and non-burst epochsshown as violin plots with individual pair values overlaid. (B) Pairwise differences in coherence (C) and dwPLI (D) across contact pairs with lines connecting burst and non-burst values.

### 2.2. Cortico-STN beta coupling during non-burst epochs does not exceed surrogate expectations

To assess whether cortico-STN beta coherence persists outside burst epochs to a degree exceeding that expected from shared spectral structure, observed coherence during non-burst periods was compared against surrogate null distributions. For each STN-cortical contact pair, up to 100 non-burst segments were randomly sampled, and surrogate datasets were generated by circularly time-shifting the STN beta signal relative to cortex (random shifts uniformly drawn between 0.5 and 5 s; n = 200 surrogates). Surrogate coherences were computed using the same procedures as the empirical data, and empirical p-values were defined as the proportion of surrogate means exceeding observed coherence. Non-burst coherence tracked surrogate expectations across recording pairs (Spearman ρ = 0.491, p < 0.001; Figure 2A), with little evidence for systematic deviation. In contrast, coherence during burst epochs showed a clear separation from surrogate distributions, as reflected by a strong leftward shift in the empirical cumulative distribution function (ECDF) of surrogate test p-values for burst epochs but not for non-burst periods (Figure 2B). The same analysis was repeated using the dwPLI values. Non-burst dwPLI closely tracked surrogate expectations (Spearman ρ = 0.703, p < 0.001; Figure 2C), whereas burst dwPLI showed separation from surrogate distributions, reflected by a leftward shift in the ECDF of burst-versus-surrogate p-values but not for non-burst periods (Figure 2D). Together, these findings indicate that cortico-STN beta coupling is predominantly confined to burst epochs.

**Figure 2.**
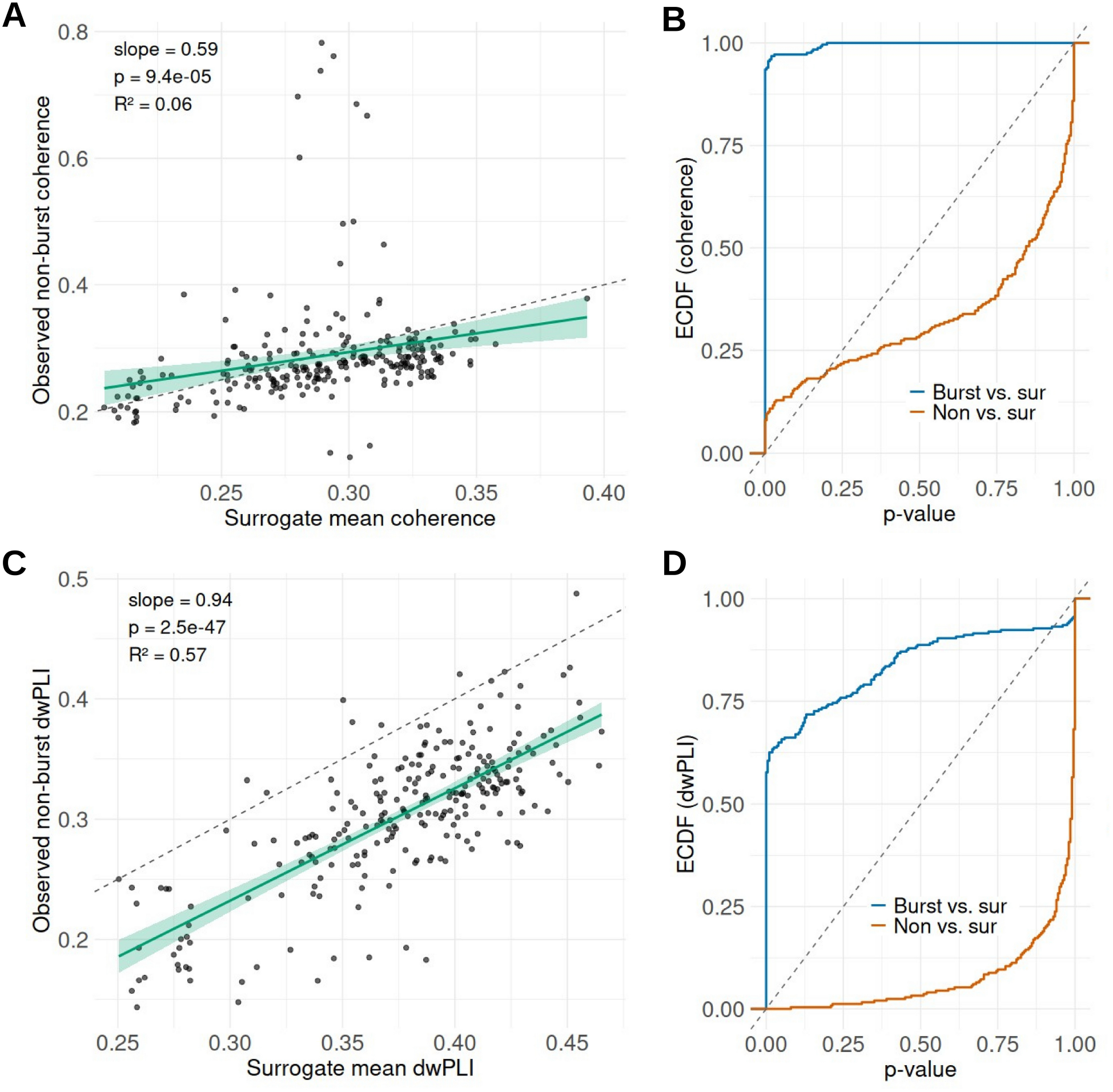
Non-burst cortico-STN beta coupling relative to surrogate control (A) Observed non-burst coherence (A) and dwPLI (C) plotted against mean surrogate coherence for each STN-cortical contact pair. Empirical cumulative distribution functions (ECDFs) of surrogate test p-values for burst and non-burst coherence (B) and dwPLI (D).

A subset of STN-cortical pairs exhibited elevated coherence during both burst and non-burst periods, with non-burst coherence exceeding surrogate expectations. These pairs were concentrated at cortical contacts near the primary motor representation, particularly contacts 9 and 10, which lay above the identity line relating observed and surrogate non-burst coherence. Quantification across contacts confirmed that a higher fraction of STN-cortical contact pairs exceeded surrogate expectations at these contacts compared with other cortical sites (Figure 3A), whereas such effects were rare or absent elsewhere. Elevated non-burst coherence at contacts 9 and 10 was observed in 6/7 and 3/7 subjects, respectively, indicating that the effect was not driven by a single individual. Importantly, despite this localized elevation in absolute coherence, burst-related increases in coherence were preserved across all contacts, with mean burst coherence exceeding mean non-burst coherence even at contacts 9 and 10 (Figure 3B). When quantified using dwPLI, this contact-specific elevation of baseline coupling was substantially reduced (Figure 3C), with fewer non-burst pairs exceeding surrogate expectations and a more uniform distribution across cortical contacts. Nevertheless, burst-related increases in dwPLI were preserved across contacts (Figure 3D). Together, these observations indicate that while a small number of motor cortical contacts exhibit stable background coupling, this does not account for the population-level effect, and pathological cortico-STN beta coupling remains preferentially expressed during burst epochs.

**Figure 3.**
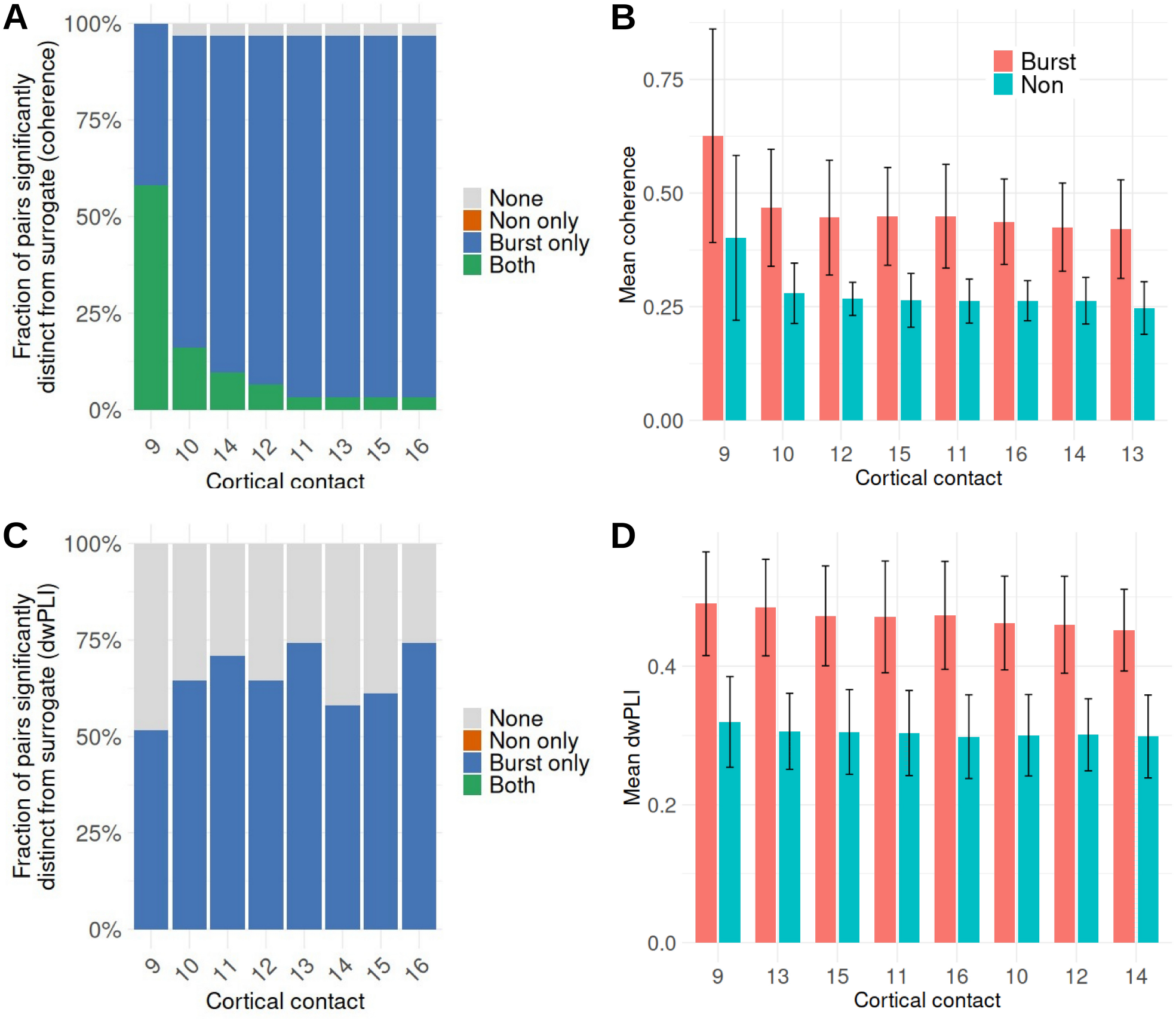
Contact-specific elevation of baseline cortico-STN beta coherence Fraction of STN-cortical pairs per cortical contact whose non-burst coherence (A) and dwPLI (C) exceed surrogate distributions. Mean burst and non-burst coherence (B) and dwPLI (D) per cortical contact. “None” indicates pairs for which neither burst nor non-burst values significantly exceeded surrogate expectations.

To test whether cortico-subthalamic beta coupling persists outside burst epochs, coherence and dwPLI during beta bursts were compared with values computed over immediately adjacent non-burst epochs of equal duration. For each STN-cortical contact pair, coupling was calculated over contiguous burst segments and over the immediately preceding and following non-burst segments matched in length to each burst. Across contact pairs, coherence and dwPLI were consistently elevated during bursts relative to both pre-burst and post-burst epochs, while pre- and post-burst values clustered closely together (Figure 4A-B). Correspondingly, the distributions of burst-pre and burst-post differences showed robust positive shifts for both metrics (Figure 4C-D), indicating that the effect was consistent across pairs rather than driven by a small subset. In contrast, pre- and post-burst values lay near the identity line across pairs (Figure 4E-F). Paired statistical analysis confirmed significantly higher coupling during bursts compared with both pre-burst and post-burst epochs (Wilcoxon signed-rank tests, all p < 2.2 × 10^-16^), with large effect sizes (coherence: paired Cohen’s d_x_ = 1.54 and 1.64 for burst-pre and burst-post, respectively; dwPLI: d_x_ = 2.06 and 2.09), whereas no meaningful difference was observed between pre- and post-burst epochs (coherence: p = 0.28, d_x_ = 0.12; dwPLI: p = 3.3 × 10^-5^, d_x_ = -0.25, mean difference ≈ -0.003). Together, these results indicate that pathological cortico-STN beta coupling is temporally constrained and does not persist immediately before or after bursts.

**Figure 4.**
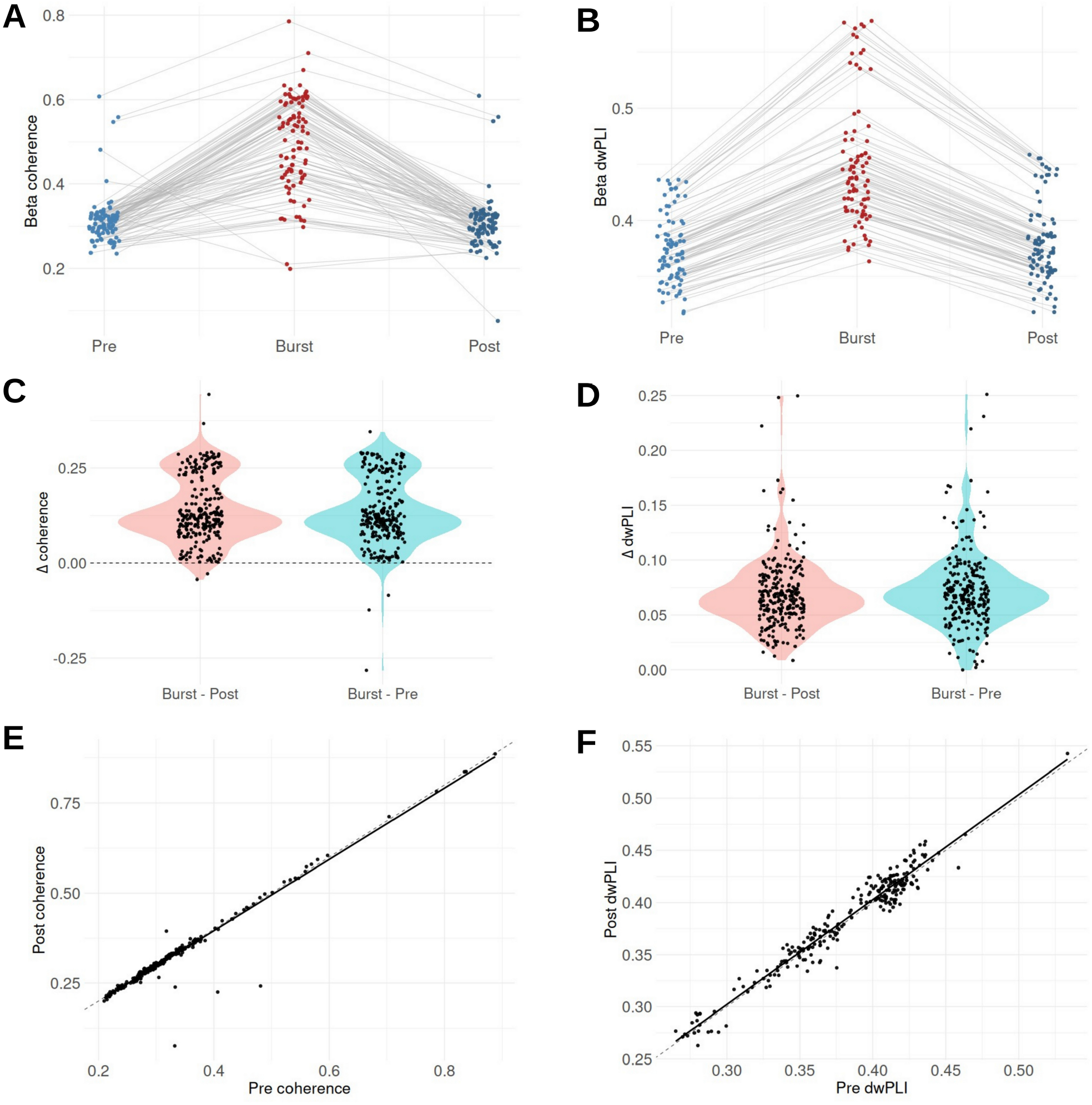
Burst-adjacent cortico-STN beta coupling Mean beta coherence (A) and dwPLI (B) during bursts and during matched pre-burst and post-burst non-burst epochs. Distributions of coherence (C) and dwPLI (D) differences for burst versus pre-burst and burst versus post-burst comparisons. Comparison of pre-burst and post-burst coherence (E) and dwPLI (F) across contact pairs.

## Discussion

This study demonstrates that pathological cortico-STN beta coupling in Parkinson’s disease is largely restricted to discrete beta burst epochs. Across burst versus non-burst comparisons, surrogate-controlled testing, and burst-adjacent analyses, beta-band coupling (coherence and dwPLI) most often collapsed to chance levels outside bursts. These findings indicate that abnormal cortico-STN synchronization is a transient phenomenon rather than a tonic property of the cortico-basal ganglia network.

Recent work has highlighted state-dependent changes in cortico-basal ganglia functional connectivity across movement and dopamine conditions, emphasizing the dynamic nature of basal ganglia network interactions rather than sustained pathological signaling[13,14]. The present findings refine this framework by demonstrating that beta-band cortico-STN coupling is temporally constrained to discrete beta burst epochs. The burst-resolved analyses reveal that measurable beta coupling collapses to surrogate levels outside bursts. Replication of these effects using dwPLI indicates that burst-specific coupling reflects consistent phase synchronization rather than purely magnitude-dependent effects.

A subset of cortico-STN pairs exhibited elevated baseline coherence during both burst and non-burst periods, localized to cortical contacts near the primary motor representation. These cases were spatially localized and did not alter the dissociation between burst and non-burst coherence. Burst-related increases in coherence remained robust at these contacts, indicating that stable background coupling does not account for the burst-specific effect. When quantified using dwPLI, this baseline elevation was reduced, suggesting that part of the non-burst coherence may reflect amplitude-dependent or zero-lag contributions rather than sustained pathological synchronization. These findings are consistent with anatomically localized baseline connectivity, possibly reflecting hyperdirect pathway interactions, distinct from transient, burst-defined coupling in PD[15–17].

The temporal confinement of pathological beta coupling has direct implications for therapeutic strategies[18]. Adaptive deep brain stimulation approaches aim to target pathological network states while sparing physiological activity. These results support the use of beta bursts and associated cortico-STN coupling as a temporally precise biomarker of pathological coupling, rather than relying on time-averaged beta power or coherence measures that conflate burst and non-burst states.

## Methods

### Participants and recordings

Intraoperative recordings were obtained from a publicly available dataset accessed through the Data Archive for the Brain Initiative (DABI; https://dabi.loni.usc.edu) under project code 1U01NS098961[19,20]. The dataset comprises recordings from seven patients undergoing unilateral STN deep brain stimulation implantation for Parkinson’s disease (sample IDs: lS01, lS02, lS03, lS04, lS05, lS06, bS20). Bipolar STN LFPs and simultaneous cortical ECoG were recorded from sensorimotor strip electrodes positioned over the perirolandic cortex. Analyses focused on STN contacts 1-8 for the right implantations and contacts 5-8 for left implantations, and on cortical contacts 9-16, corresponding to approximate primary motor (contacts 9-12) and primary somatosensory cortex (contacts 13-16), based on intraoperative electrode placement spanning the perirolandic (central sulcus) region.

### Preprocessing and beta burst detection

All signal processing was performed using MATLAB (MathWorks; version R2025B). Raw signals were high-pass filtered at 1 Hz and band-pass filtered in the beta range (13-30 Hz) using zero-phase fourth-order IIR filters designed with designfilt and applied using filtfilt. To minimize edge artifacts, 2 s were trimmed from the beginning and end of each filtered signal, and the first 60 s were discarded. Sampling frequency was extracted from metadata or set to 1000 Hz if unavailable. Beta bursts were algorithmically identified from the STN beta-band signal. The analytic amplitude envelope was computed using the Hilbert transform (hilbert) and smoothed with a 50 ms moving average (smoothdata, moving mean). Burst epochs were defined as contiguous samples exceeding the 75th percentile of the smoothed envelope for each contact, while non-burst epochs were defined as samples below the 50th percentile[8]. The 50th percentile threshold was selected to ensure that non-burst epochs reflected subthreshold beta activity while avoiding transitional periods near the burst boundary. Contiguous segments shorter than 100 ms were excluded. One STN-cortical contact pair exhibited only a single detected beta burst and was excluded from burst-adjacent analyses.

### Coherence and phase-based coupling estimation

Cortico-STN beta coherence was estimated using the MATLAB mscohere function (Welch’s method)[21]. For each STN-cortical contact pair, coherence was computed separately for burst and non-burst epochs and averaged across frequency bins within the beta band. Beta frequency indices were defined from a representative burst segment to ensure consistent frequency averaging across conditions. To avoid bias due to unequal numbers of non-burst segments, up to 100 non-burst segments per contact pair were randomly sampled for coupling estimation. The debiased weighted phase lag index (dwPLI) was calculated from the imaginary component of the cross-spectrum to quantify non-zero-lag phase synchronization, following Vinck *et al*.*[22]*. Coherence captures coupling magnitude, whereas dwPLI quantifies non-zero-lag phase synchronization and reduces sensitivity to shared amplitude or volume conduction. Burst and non-burst dwPLI values were computed over the same segments used for coherence analysis. Mean coherence and dwPLI values were computed for each condition, yielding per-pair estimates of burst coupling, non-burst coupling, and their difference.

### Surrogate null testing

To determine whether coupling (coherence and dwPLI) observed during non-burst periods exceeded that expected from shared spectral structure or finite sample effects, surrogate null distributions were generated using circular time-shifting[23]. For each STN-cortical contact pair, the STN beta-band signal was circularly shifted relative to the cortical signal by a random offset uniformly drawn between 0.5 and 5 s, preserving spectral content while disrupting temporal alignment. By default, 200 surrogate datasets were generated per test. Surrogate coupling values were computed using the same procedures as for the empirical data. Empirical p-values were defined as the proportion of surrogate mean coupling values greater than or equal to the observed mean coupling. Separate surrogate tests were performed for burst and non-burst epochs. Random number generation was seeded deterministically (seed = 1) to ensure reproducibility.

### Time-resolved burst-aligned coherence analysis

To test whether cortico-STN beta coupling persisted immediately before or after bursts, a time-resolved, burst-aligned analysis was performed. Coupling metrics were computed in sliding windows centered on burst onsets over a peri-burst interval extending 50 ms before to 50 ms after burst onset. Window length (15 ms) and step size (5 ms) were specified a priori, and coupling within each window was computed with the MATLAB mscohere (for coherence) and the dwPLI implementation described above (for phase-based coupling) using the chosen window and 50% overlap. For comparison, matched non-burst windows of identical duration were sampled from time points at least 50 ms away from any burst, ensuring temporal separation from burst-related activity. Coupling traces were averaged across bursts for each contact pair to yield mean burst-aligned and matched non-burst time courses.

### Statistical analysis

Paired comparisons between burst and non-burst coupling values were assessed using Wilcoxon signed-rank tests, and paired t-tests are reported for completeness[24]. Effect sizes were quantified using paired Cohen’s d[25]. Correlations between observed and surrogate coupling values were assessed using Spearman’s rank correlation coefficient. Linear mixed-effects models with random intercepts for patient and electrode contact were fit in R (version 4.5.1) using the lme4 and lmerTest packages[26]. All statistical tests were two-sided unless otherwise stated.

## Acknowledgements

The research was supported by the National Institute for Health Research (NIHR) Oxford Biomedical Research Centre (BRC). The views expressed are those of the author(s) and not necessarily those of the NHS, the NIHR or the Department of Health.

## Data availability

All MATLAB and R analysis scripts are available at https://github.com/cabeaudoin/beta_burst_nonburst.git.

## Notes

https://github.com/cabeaudoin/beta_burst_nonburst.git

### Competing Interest Statement

The authors have declared no competing interest.

